# Stage specific classification of DEGs via statistical profiling and network analysis reveals potential biomarker associated with various stages of TB

**DOI:** 10.1101/414110

**Authors:** Romana Ishrat

**Affiliations:** CIRBSs, JMI University, New Delhi-110025, India

**Keywords:** LTTB, DEGs, Gene Ontology; KEGG, network analysis, gene knock-out, LCP

## Abstract

**Background:** Tuberculosis (TB) is a deadly transmissible disease that can infect almost any body-part of the host but is mostly infect the lungs. It is one of the top 10 causes of death worldwide. In the 30 high TB burden countries, 87% of new TB cases occurred in 2016. Seven countries: India, Indonesia, China, Philippines, Pakistan, Nigeria, and South Africa accounted for 64% of the new TB cases. To stop the infection and progression of the disease, early detection of TB is important. In our study, we used microarray data set and compared the gene expression profiles obtained from blood samples of patients with different datasets of Healthy control, Latent infection, Active TB and performed network-based analysis of DEGs to identify potential biomarker.

**Objectives:** We want to observe the transition of genes from normal condition to different stages of the TB and identify, annotate those genes/pathways/processes that play key role in the progression of TB disease during its cyclic interventions in human body.

**Results:** We identified 319 genes that are differentially expressed in various stages of TB (Normal to LTTB, Normal to Active TB and LTTB to active TB) and allocated to pathways from multiple databases which comprised of curated class of associated genes. These pathway’s importance was then evaluated according to the no. of DEGs present in the pathway and these genes show the broad spectrum of processes that take part in every state. In addition, we studied the regulatory networks of these classified genes, network analysis does consider the interactions between genes (specific for TB) or proteins provide us new facts about TB disease, which in turn can be used for potential biomarkers identification. We identified total 29 biomarkers from various comparison groups of TB stages in which 14 genes are over expressed as host responses against pathogen, but 15 genes are down regulated that means these genes has allowed the process of host defense to cease and give time to pathogen for its progression.

**Conclusions:** This study revealed that gene-expression profiles can be used to identify and classified the genes on stage specific pattern among normal, LTTB and active TB and network modules associated with various stages of TB were elucidated, which in turn provided a basis for the identification of potential pathways and key regulatory genes that may be involved in progression of TB disease.

## Introduction

TB is a communicable disease generally caused by the bacterium Mycobacterium tuberculosis (MTB). The lungs are typically affected (pulmonary TB) but other body parts can be also affected (extra pulmonary TB)[1]. The disease spread through air when a person who are infected with active TB expel out bacteria, for example by coughing and sneezing[2]. In 2016, 10.4**M** individuals infected with TB disease and 1.7**M** died from the disease (including 0.4 million among people with HIV, 40% of HIV deaths were due to TB). TB kills more adults in India than any other infectious disease (in 2016 an estimated 28 lakh cases occurred and 4.5 lakh people died due to TB). India has the highest burden of both TB and advanced TB (like MDR TB) and second highest of HIV associated TB. In India, the major challenges to curb the TB are poor primary healthcare system in rural areas; due to deregulation of private health care leading to indiscriminate use of I^st^ & II^nd^-line TB drugs; poverty; spreading HIV-infection; lack of administrative coordination among government functionaries bodies. In our current study, we used meta-analysis of individual raw microarray data (GSE series) deposited in GEO database, got from blood samples of individuals with different datasets (e.g., Controls vs. TB disease, Control vs. Latent TB, Latent TB vs. TB disease). For subsequent analysis, we were selected that showed a significant differential expression in most of experiments. We have performed gene-transition study from DEGs data and text mining between different stages of TB and classified the DEGs in stage specific manner like Normal to Latent TB infection, Normal to Active TB and Latent TB infection to Active TB, then identified specific interaction network modules and figure out its topological properties to anticipate key regulators among which few have fundamental significance for their biological activities and regulating mechanism. The present method to prioritize the disease genes is mainly centred on the ‘guilt-by-association’ presumption, that means the physically and functionally linked genes are possibly participated in the same biological-pathways having comparable effects on the phenotypes[3]. The concept of network theory is an imperative method to know the topological properties and the complex-systems dynamics correspond to their functional modules. The complex networks may be classified into four types of networks: (a) scale-free network, (b) small-world network, (c) random network and (d) hierarchical network. For the biologist, hierarchical network has special interest because it incorporates the mien of modules and distributed hubs (sparsely) regulate the network. In our study, the goal was to give diagnostics a way to define the stages of infection to make specific remediation.

## Results

### Differentially Expressed Genes (DEGs)

A total of 5680 DEGs were identified after the extensive analysis of all the 11 GSE series; of which 2660 were up regulated and 3020 were down regulated genes. The DEGs were divided into five different comparison groups, Normal vs. Latent Infection, Normal vs. active TB and Latent infection vs. Active TB.

### Classification and Overrepresentation Analysis

A total of 5,680 differentially expressed genes were clustered according to ‘GO-MF (Molecular Function)’, ‘GO-BP (Biological Process)’ and ‘PANTHER Protein Class’ shown in (Figure 1). All these predictive DEGs show a broad spectrum of protein classes involved in a wide array of processes like binding proteins (RNA and DNA), helicases and nucleases are found within the “Nucleic Acid Binding” protein-class. The “Enzyme Modulator” category features kinase, G-protein, phosphatase and protease-modulators. Structural motif and nuclear hormone receptors are part of the “Transcription Factor” protein class. The “Hydrolases” is a sub-category of Proteases and Phosphatases. The “Receptor” protein class includes cytokine-receptors, protein-kinase receptors, ligand-gated ion channels, nuclear-hormone receptors and G protein coupled receptors. Besides these proteins classes, signaling molecules, Transferase, oxidoreductase and transporter are most abundant protein classes. The two most abundant GO-Biological Process groups—“Metabolic Process” and “Cellular Process” which is not surprising as these contains genes involved in the most basic life processes. The “Cellular Process” includes cell cycle, cell-cell signaling, cell component movement, growth and proliferation and cytokinesis. The heading “Metabolic Process” includes carbohydrate metabolism, cellular amino acid metabolism, lipid metabolism, nucleobase-containing compound metabolism, protein metabolism and the tricarboxylic acid cycle. “Biological Regulation” includes metabolism, cell cycle, the regulation of apoptosis, catalytic activity, translation and homeostasis. Beside all these biological processes response to stimulus, localization and developmental process also represent the significant number of protein. To know the probability of largely occupied protein classes and GO category among the DEGs, we used PANTHER’s overrepresentation analysis. When we compared with the reference genome, we found that any of the most abundant categories are overrepresented in the data (Table_1). The five most abundant protein classes “Chemokine”, “Cytokine”, “Hydrolase”, “Ribosomal protein” and “RNA binding protein” were enriched along with the classes “Signaling molecule” and “Cell adhesion molecule”. The five highly populated GO-Biological Processes were also enriched in “Cytokine-mediated signaling pathway”, “Response to external stimulus”, “Immune response”, “Locomotion”, and “Signal transduction” besides these major classes “Cell communication”, “Developmental process”, “Cellular process” and “MAPK cascade” were also enriched in GO Biological Processes. Finally, the top six GO Molecular Functions that were enriched are “Chemokine activity”, “Cytokine activity”, “Cytokine receptor binding”, “Oxidoreductase activity”, and “Receptor binding”.

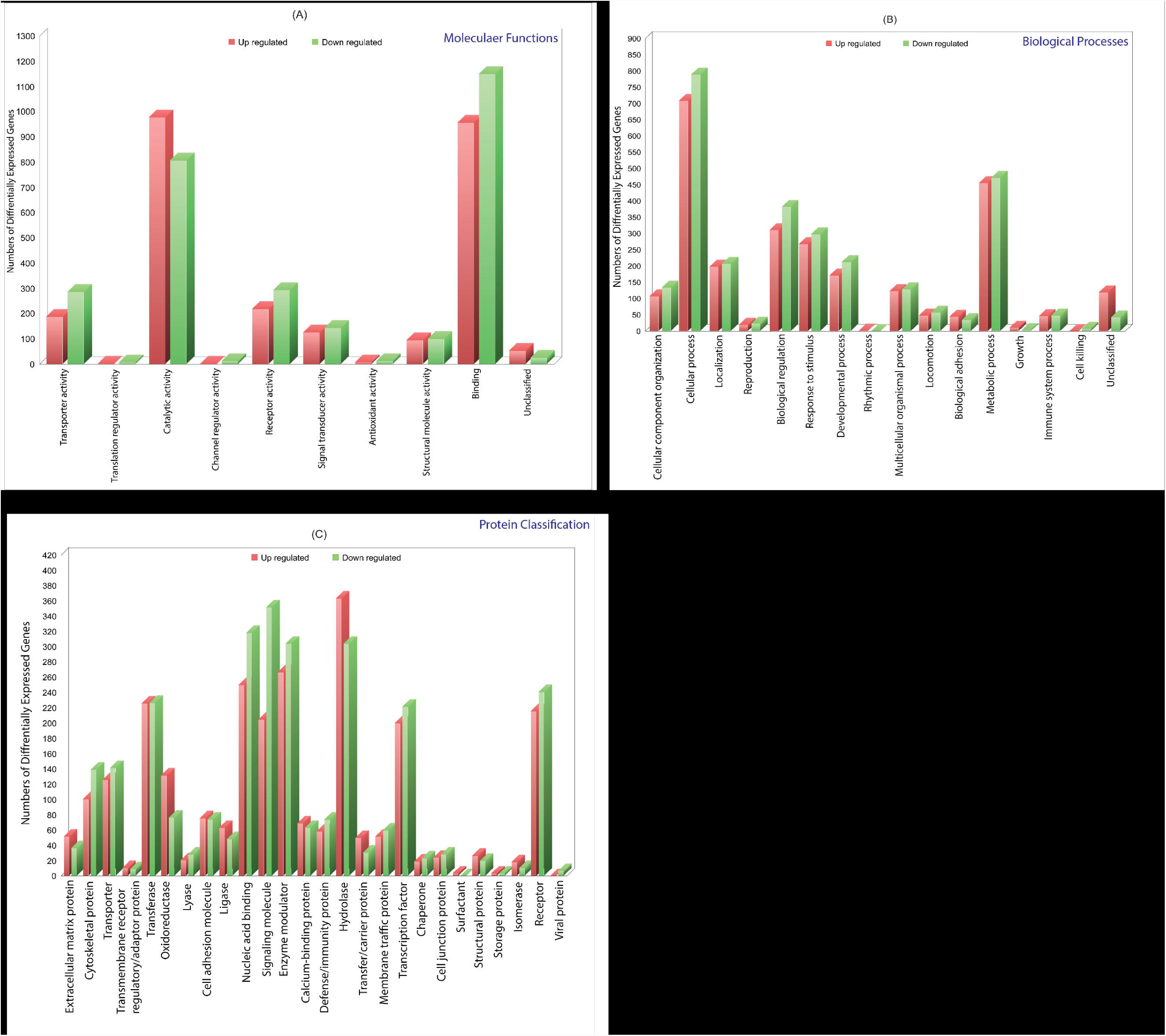

### Gene-Transition and GO Enrichment Analysis

We filtered out genes that are common in more than one series of data for further study (Table_2). A GO term enrichment analysis was performed to gain a deeper knowledge of these DEGs. We isolated up and down regulated genes from each stage and performed GO Enrichment analysis using DAVID tool. The molecular pathways associated with the differentially expressed genes were identified using KEGG pathway analysis (Table_3 and 4).

#### • Normal to Latent infection

In this case, we found 12 genes (*IER5L, MS4A6A, DOK2, FZD2, NCKI-ASI, SNHG12, NLRC4, XPO7, SMA4, CD36, AFFI* and *NDUFS8*) are up regulated and 16 genes (IL1A, *IL6, ACOD1, IL1B, ELOVL7, PTGS2, EREG, F3, IFIT1, TNF, KANK1, CCL4, CXCL11, PTX3, IRAK2* and *AREG*) are down regulated. The GO analysis of these DEGs are enriched in “Immune response”, “Inflammatory response”, “Chemokine activity”, “Cellular process”, “Biological regulation”, “Regulation of metabolic process”, “Protein binding”, “Catalytic activity”, “Bindings” etc. On pathway analysis, most of the DEGs are down regulated (beneficial to pathogen) that are enriched in very important pathways like Toll-like receptors, NF-Kappa B signaling, Cytokin-Cytokin receptor interaction, MAPK signaling pathway, Tuberculosis and TNF signaling.

#### • Normal to Active TB disease

We found total 153 genes (UP regulated) and 119 genes (Down regulated) among the various cases. In this stage, the DEGs were not only enriched in “Inflammatory response”, “Immune response”, “Signal transduction”, “Response to stimulas”, “Apoptotic process”, “Cellular process”, “Metabolic process”, “Binding”, “Catalytic activity”, “Receptor activity” etc. and the most abundant enriched pathways (for up regulated genes) in this stage belonged to “Cytokine-cytokine receptor interaction “Chemokine signaling”,Toll-like receptor signaling pathway”, “NF-kappa B signaling pathway”, “Transcriptional misregulation in cancer”, “Pathways in cancer” and for down regulated genes, the enriched pathways are “Cell cycle”, “Chemokine signaling pathway”, “NF-kappa B signaling pathway”, “TNF signaling pathway”, “PI3K-Akt signaling pathway, “Metabolic pathways” and “MAPK signaling pathway”.

#### • Latent infection to Active disease

In this case, we found that 154 genes are up regulated and 84 genes are down regulated. Compared with healthy controls, TB-disease appeared to be mostly related to “Binding”, “Catalytic activity”, “Receptor activity”, “Transporter activity”, “Cellular and metabolic process”, “Biological regulation”, “Immune response”, “Oxidoreductase activity” etc. and the most abundant enriched pathway (for up regulated genes) in this stage belonged to “Metabolic Pathway”, “Nod like receptor signaling pathway”, “Regulation of Actin cytoskeleton”, “Biosynthesis of antibiotics”, “Metabolism of xenobiotics by cytochrome p450”, “Thyroid hormone synthesis”, “Regulation of autophagy”, “Peroxisome” and for down regulated genes, the enriched pathways are “Chemokine signaling pathway”, “TNF signaling pathway”, “RAP1 signaling pathway”, “PI3K_AKT signaling”, “Metabolic process”, “Apoptotic”, “Focal Adhesion”, “MAPK signaling pathway”, “MicroRNAs in cancer” and “Amoebiases”.

### Gene interaction: Hierarchical Scale-free Network

The classified genes of various stages of TB (**Table_2**) were used to construct their regulatory network. We have constructed six networks for UP and Down regulated genes separately. The topological parameters of the network obey power law distributions (as a function of degree). The probability of clustering co-efficient *C(k)*, degree distributions *P(k)*, and neighborhood connectivity *C*_*N*_*(k)* exhibit power law or fractal nature shown in **Figure 2**. The results for the all the networks are summarized as follows,

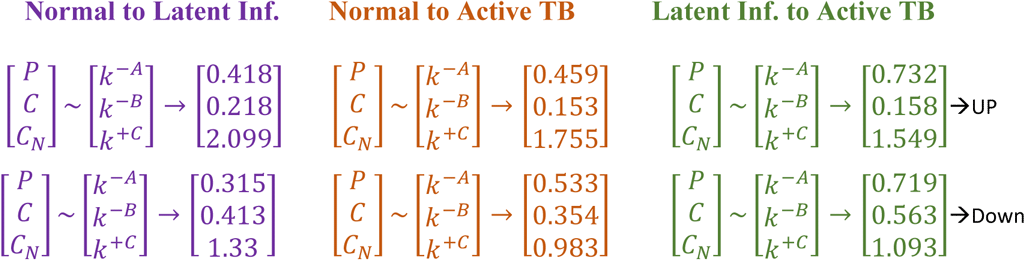

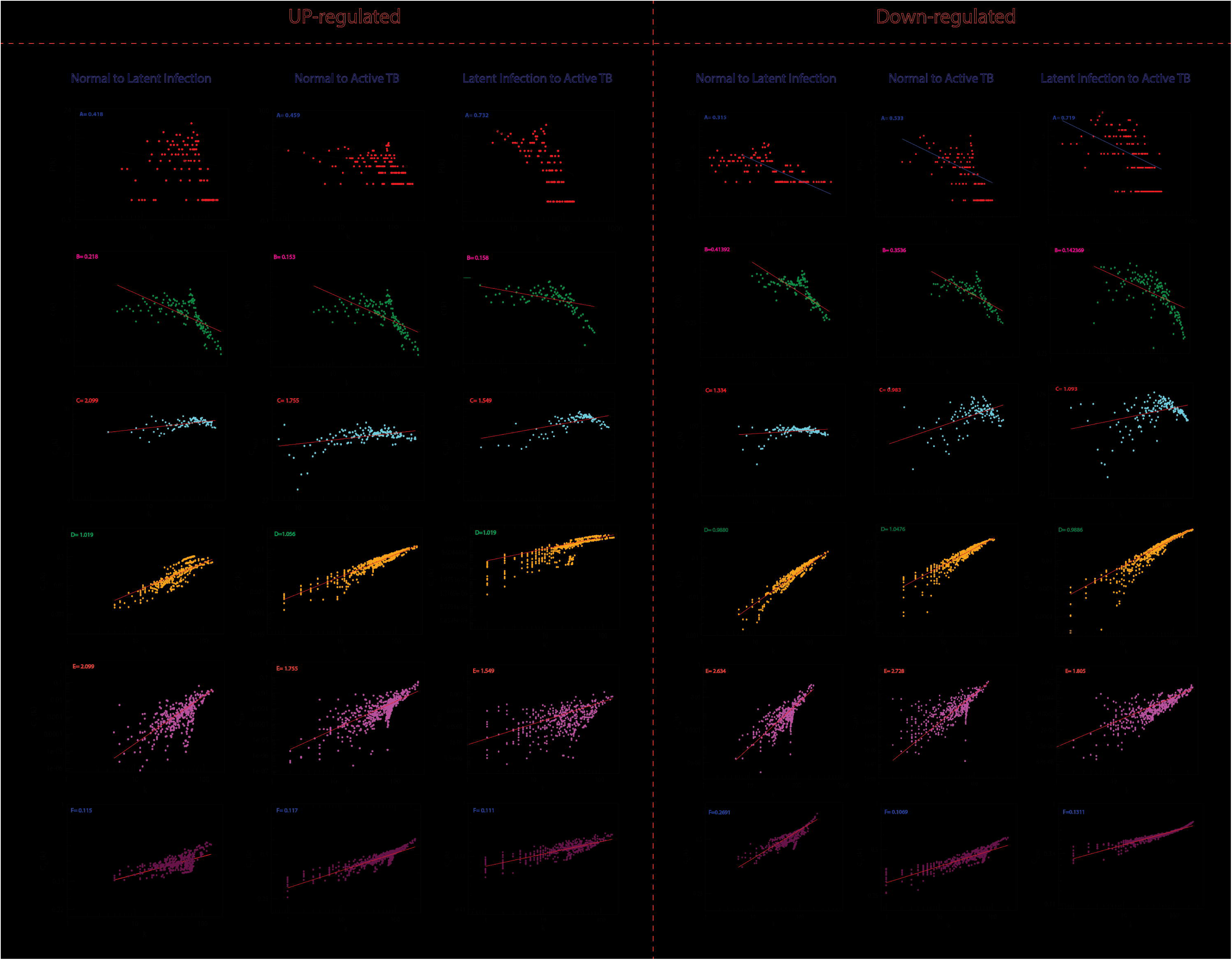

This network behaviour indicates hierarchical scale free network[4] [5] [6]. The power law fits on the data points of the network’s topological parameters are done and confirmed by following the standard statistical fitting method given by Clauset et al[7]. where, the p values to all data sets were calculated (against 2500 random samplings) that is greater than 0.1 and data fitting goodness is less. **P(k)** and **C(k)** have negative values that mean the network follows hierarchical pattern and positive-value of **C_N_(k)** that means the network follow the assortativity that identifies a huge cluster of degree-nodes (rich club) which regulates the network. The centrality parameters: betweenness (C_B_), closeness (C_C_) and eigenvector (C_E_) centralities of the network also show fractal behaviour and good connectivity of nodes in a network is distinguished by eigenvector or centrality C_E_(k). It calculates the effectiveness of the spreading (receiving) power of data of nodes from the network. These properties follow the power law behaviours as follows,

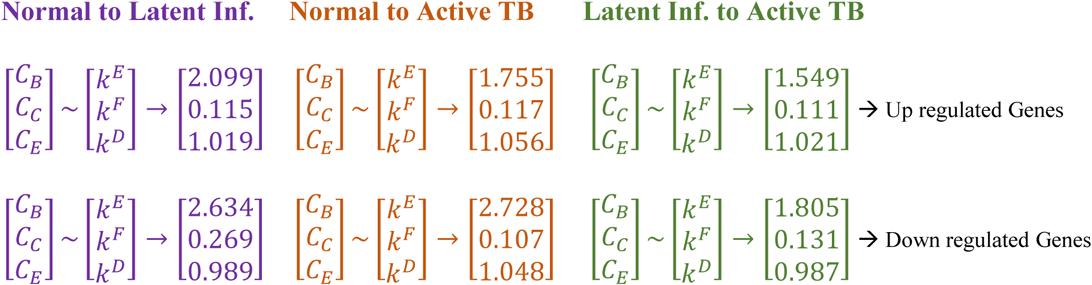

### Identification of key regulators and properties

Since the popularity of leading hubs get change according to gene activities and its regulation, so we can’t say that all the predictive leading hubs are key regulators for disease progression but few of these hubs can play significant role, which we called them as fundamental key regulators (FKR). The structure of modular and its arrangement are done through Newman and Girvan’s standard community finding algorithm at various levels of the organisation. Using this community finding algorithm, we found that our six networks are hierarchically organised at various levels (**Suppli_1A,B**). The Hamiltonian Energy (HE) and corresponding modularity (QN) (as a function of levels of organization) are found to be decreased as one goes from top to down organisation (**Figure 3**).

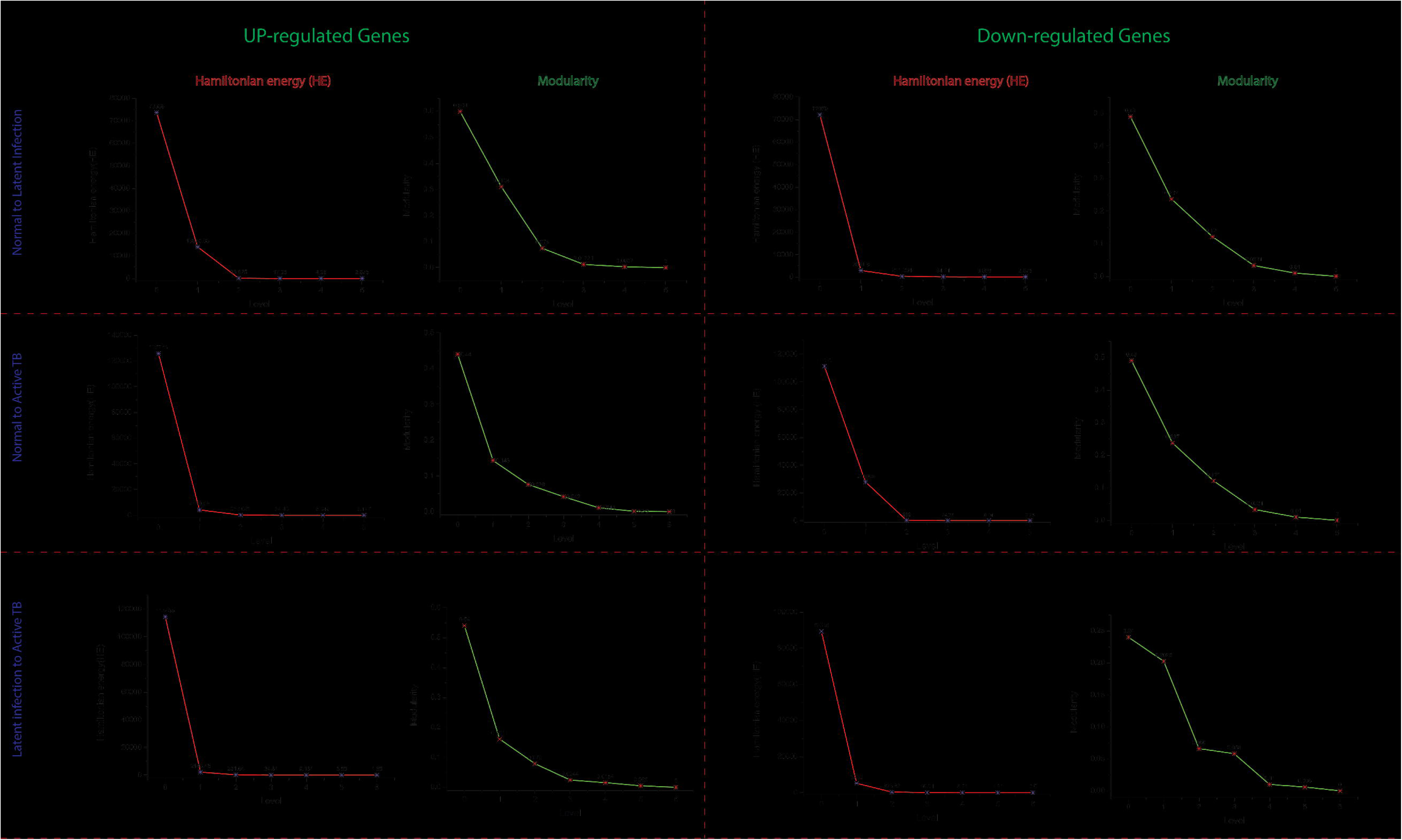

Following the definition of KR, we identified total 29 key regulators from various stages of TB. In normal to latent infection, we found eight genes which are ‘FZD2, NDUFS8, NLRC4’ (Up regulated) and ‘CCL4, IL1B, IL1A, TNF, AREG’ (Down regulated), While in latent to active TB, we found 15 genes which are ‘EIF2AK2, SAMD9L, IFI44L, DDX58, IFI44, HERC5, NFE2L3, LRRK2’ (Up regulated) and ‘CSF3, MAP3K8, IRAK2, TNFAIP3, TRAF1, PLAUR, CD44’ (Down regulated) and similarly for normal to active TB disease, six genes ‘TST, MTHFD1, CARS2’ (Up regulated) and ‘ETV6, NFKB1, MET’ (Down regulated) are fundamental key regulators. Datamining using **Geneclip 2.0**[8] [9] to show that 28 of 29 genes were enriched in several biological processes shown in **Suppli-2**.

To understand the regulating ability of each of the 29 KRs, we calculated the Probability *P*_*x*_ (*y* ^*l*^).

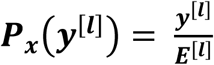

Where **x** shows the no. of edges **(*y*^*1*^])** at level **(l)** and **(*E*[^*1*^])** is total no. of edges of the network or modules and sub-modules. The measured probability *P*_*x*_ (***y*** ^***l***^) of all the KR show an increase in **Px** as level increases (top to bottom direction). At deeper level of the organisation, regulation of FKR increases and their activities become more prominent. Therefore, these FKR become backbone of the network-organization, stabilization & active workers at grassroot level (**Figure 4**).

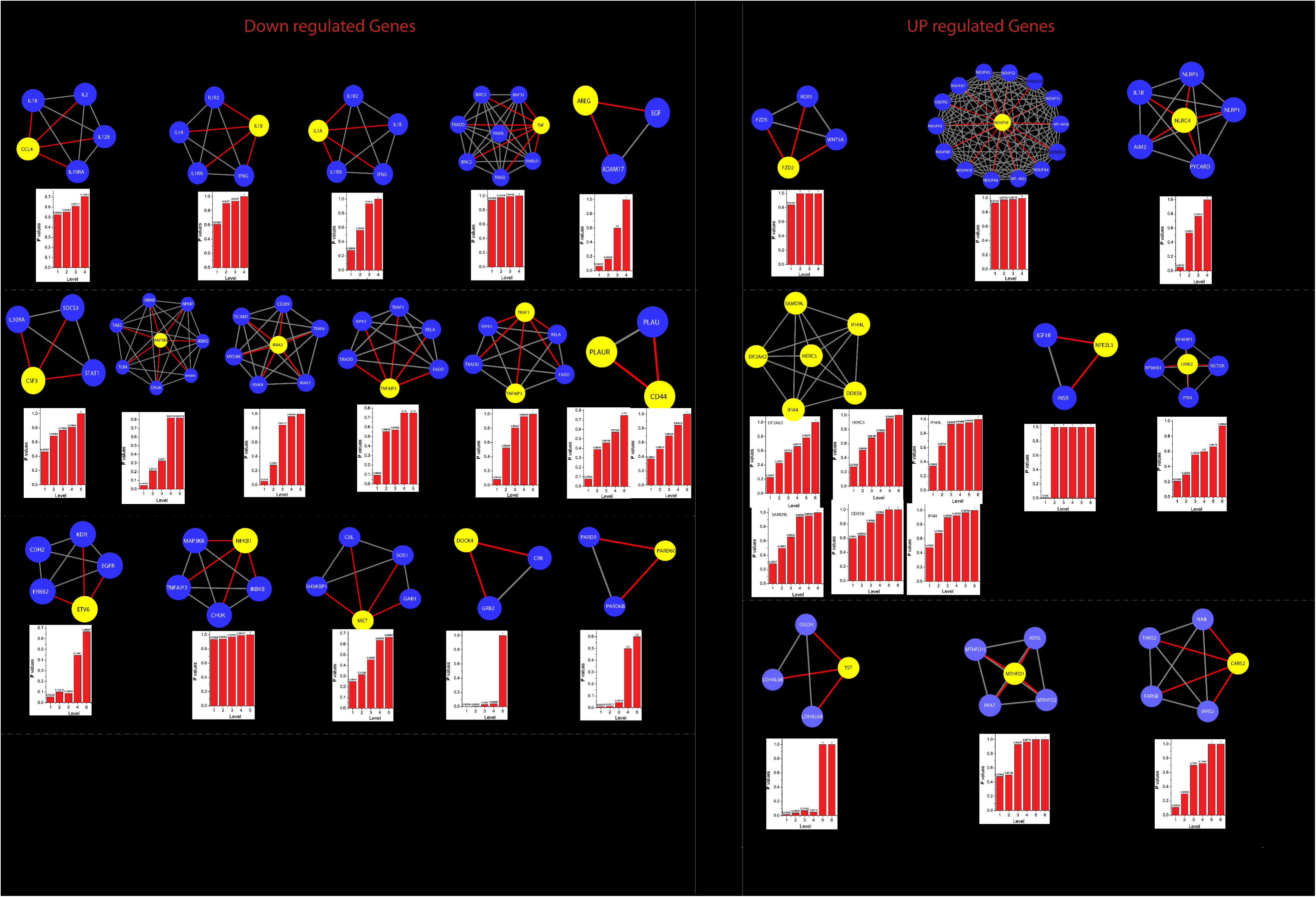

### Local perturbations driven by key regulators

The knock-out experiment of all the hubs/motif from the parent networks could able to highlight the local perturbations driven by these individual hub or motif, and their effect on global network properties. It has been revealed that the network is tolerant to hub’s deletion that means the important network-elements are still remain after elimination of hubs at level 0 (parent network). One of main cause is that the P-P interactions networks are too dense to be broken into fragments by only removing hubs. However, the elimination of these hubs/motif from the complete network does cause significant variations in the topological properties of the network, where, **P(k)** and **C(k)** change significantly in complete network level, whereas **C_N_(k)** change slightly. Likewise, the variations in the exponents of centrality measurements (ǫ, η and δ) also significant changes (**Figure_5**). Since, it is clear from the differences in the exponents of topological parameters that network perturbation increases when on goes to deeper level (top to down direction). In our case, most of the perturbation increases after the 3rd level, at this level elimination of key regulator from network almost breakdown the submodules existing in the deeper levels, such type of behaviour shows that local perturbation is highest at deeper levels.

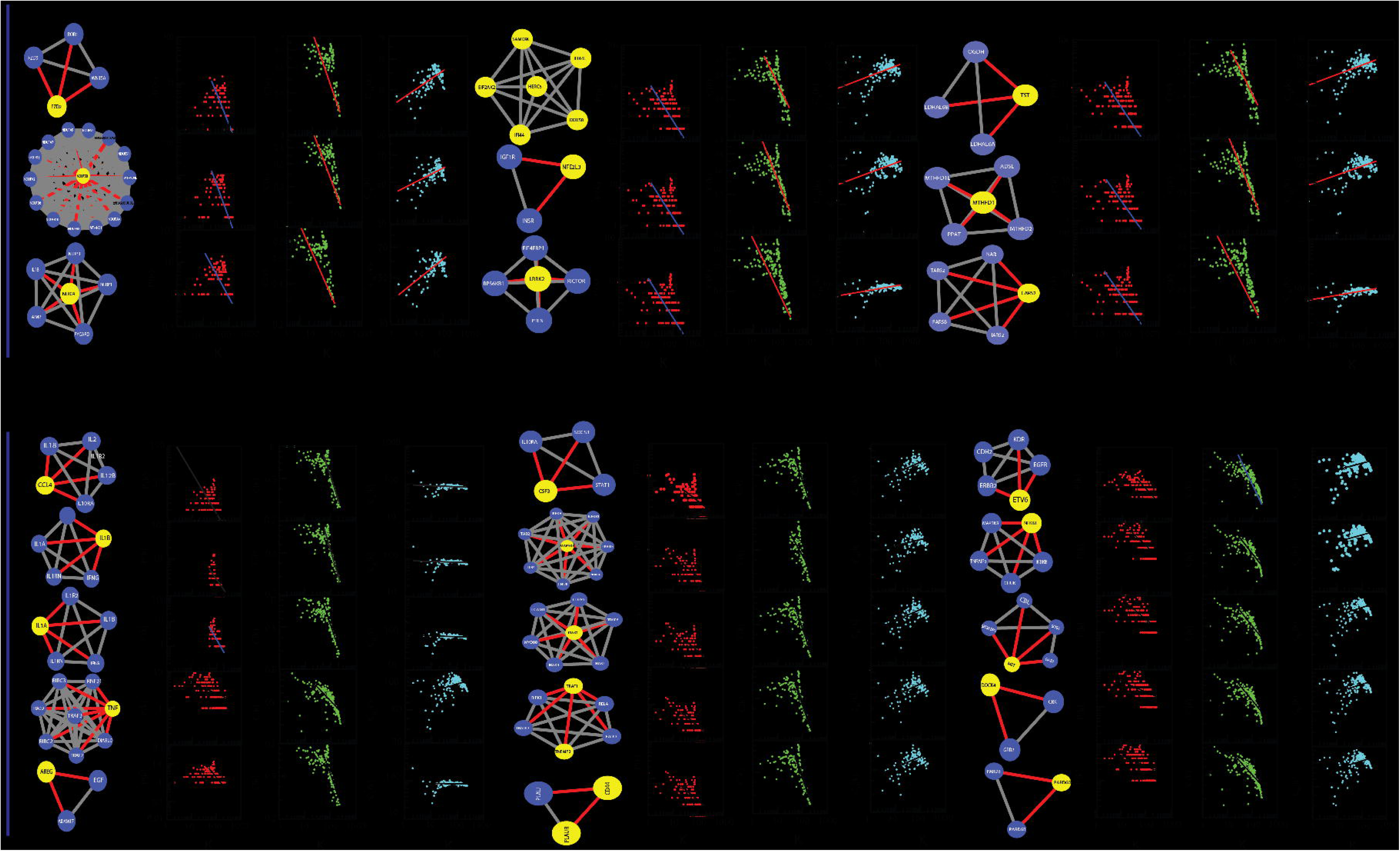

### The local-community-paradigm: Evidence of self-organization

The LCP architecture supporting the quick transfer of data across the several network-modules and through the local processing too. We have analysed all the six networks to check the maintenance of its self-organization at different levels using LCP method. For different level, the calculated LCP-corr of all the modules or sub-modules are shown in **(Figure_6)**. The average values of LCP-corr (we ignored modules which having zero LCP-corr) are greater than 0.853 at each level. This shows that the network maintains compactness, self-organisation and has efficient data processing. It characterizes robust LCP-networks that are dynamic in nature and heterogeneous, which help in network re-organization and evolution.

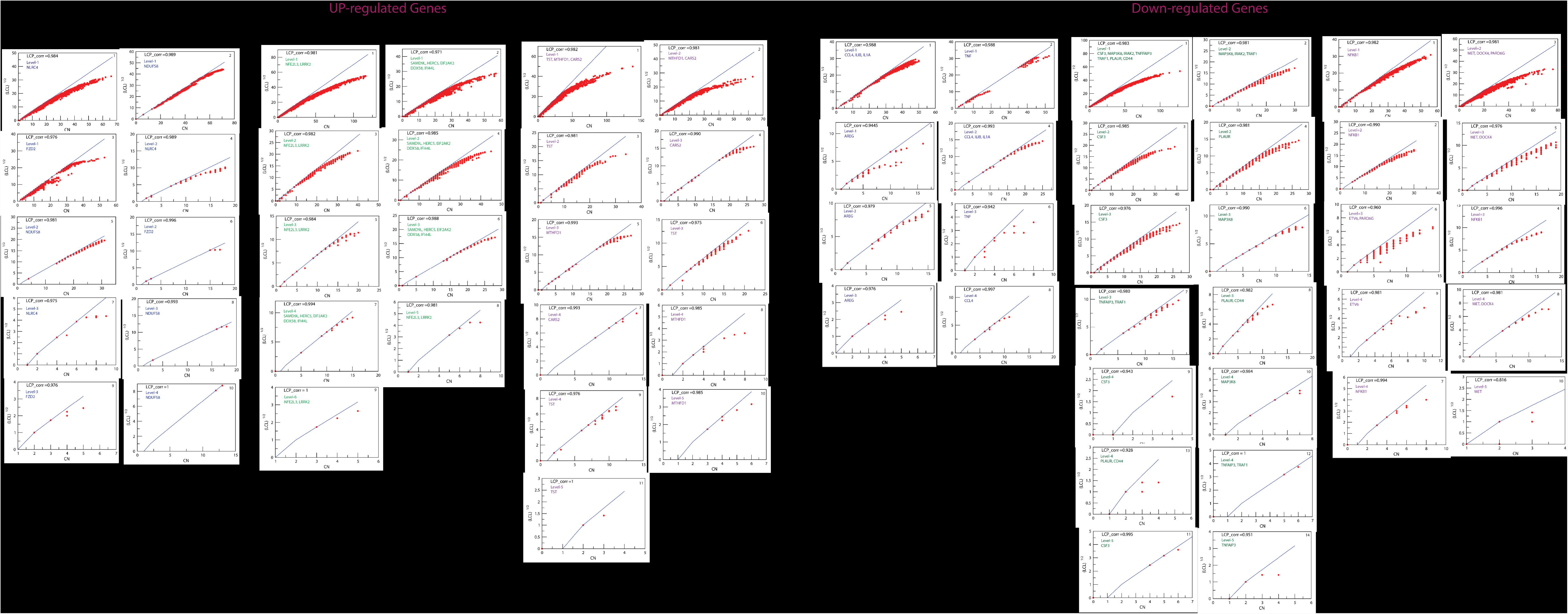

## Discussion

TB remains one of the most significant infectious disease as a leading cause of death worldwide; the present challenge is to develop a delicate & effective method to identify the latent TB infection (LTBI). Once pathogen enter into the bloodstream of host, it can infect several tissues like lung, pancreas, or thyroid and heart skeletal muscles. However, in latent TB stage, the pathogen remains in inactive form for many years before transforming into infectious form or TB-disease. However, after TB-medication, there will be a chance to reactivation of MTB due to immunosuppressant/MDR in TB bacteria[10] [11]. TB is a terrible communicable disease in that 90% of cases of latent infection with MTB not show any symptoms/sign but have a 10% lifetime possibility of transforming into active-TB. To stop the disease epidemic, early diagnosing-method or techniques are required. Gene expression profiling have uncovered the differences in the transcriptome among normal condition, latent infection and active TB. These results not only revealed significant genetic biomarkers indicative of, LTBI, TB-disease conditions but also recognized transcriptionally regulated genes that varies in biological-functions.

In our study, we found that the most of DEGs relative to normal to TB disease condition (Including Latent Infection) are enriched in important pathway like Toll-like receptors. In fact, pathogen is identified by the receptors on the surface of immune cells, and Toll-like receptors (TLRs) are one of them. Different TLRs including TLR2, 4, 9 and 8 play important roles in TB infection[12]. These receptors (Toll-like NOD like etc.) are expressed irrespective of whether they participate in immune signalling or immunity. The interaction of MTB with these receptors (like TLRs) initiates an intercellular signaling cascade that culminates in a pro-inflammatory response (cytokines and chemokines that serve as a signal for infection).

### Cascade of events going on in response to the TB infection cycle

According to the cycle of infection by Young et al., 2008, the exposure of the microorganism up to its development to latent stage, there are stringent amount of processes have appeared. The long fight of MTB to establish its generation by stabilizing its environment for metabolism have evolved with an ability to overcome immunity. In this scenario of transferring from Initial infection (say Normal) stage to Latent stage have a profound increase in the overall immune response outside the Hcell (Host cell) and an increased intensity of basic cellular machine establishment process for the Pcell (Pathogen Cell) as shown in **Table 1, 3**. In case, the elimination eliminates the eliminator even after the induction of T-cell but able to suppress the effect, Pcells have two choices, either develop army or attack the adaption of immunity. In order it decides to enter the resting phase (i.e. latent) Mtb’s survival instincts allows it to develop its metabolic environment by down regulating antigen presentation and the release of cytokines too (like IFNγ). On the other hand, if adaptive immunity has taken its chance then Mtb takes the defensive mode by subverting various normal cell cycle functioning inside the macrophages thus making a halt on the maturation of phagosomes. Mtb does this by downregulating receptor signalling pathways, cell cycle mediating molecules, disturbing metabolism also by perturbing apoptosis related gene in order to survive the acute phases (active TB).

### Events benefited to Pathogen

Taking cycle of infection into consideration it is very difficult to say which process(s) is beneficial to pathogen at which stage. As it tries to control the expression of bunch of genes that are involved in many different pathways (all at the same time) in order to overcome the pathological elimination from the host cell. Once Mtb or any other antigen enters the host it initiated a series of action in order to recognise it as self or non-self-molecules. This action is taken up by the wandering cells (Macrophages, dendritic and B cells) that recognise antigen by PRRs (pattern recognition receptors) thus, initiating the release of inflammatory mediators (PTGS2 etc.) that summons more of the macrophages to release cytokines (like TNF, IL-1 etc.) and initiate complement by TLRs to call for immediate adaptive (humoral kind) immunity. Mtb in case finds it difficult to precede it tries to down regulate all these processes to enter into the latent state as evident from the **Table 3, 4.** In case Mtb has to re-infect the host, it prepares itself by up-regulating the basic metabolic processes (like Localization, Cell adhesion, Protein phosphorylation and biological regulation) by increasing the expression of set of genes. In the meantime, it also able to recognise some antibiotics and metabolise, it is using cytochrome p450 (called Response to stimulus). Since, in the due course of its preparation processes it has established in a very hostile environment (i.e. within active immune response in host with vaccination) it does it by further down regulating another set of genes that are responsible in host cell to raise much more furious attack on Mtb.

### Some beneficial events with respect to Host cell reverted by pathogen

After breaching of the first line of defence (Mechanical barriers mucus layers) pathogens are captured by APCs (Antigen presenting cells) like dendrites and macrophages. NF-Kappa B signaling pathway is central to the host’s response to many pathogens. TNF with NF-Kappa B signaling play important role in infection dynamics in humans and multiple animal systems[13]. Cytokines released by macrophages are important intercellular regulators and mobilizers of cells to get engaged in innate as well as adaptive inflammatory host defences, cell growth, differentiation, cell death, angiogenesis, and development and repair processes aimed at the restoration of homeostasis[14]. A number of cytokines (including TNFα) are regulated by MAPK signaling pathway that are released by the macrophages infected with *MTB*[15] [16], TNF signaling if correctly activated help in providing resistance to mycobacteria by inhibiting bacterial growth and macrophage death[16]. Another set of chemicals like chemokines (a family of small cytokines) activated in response infection attract immune system cells (other leukocytes) at sites of inflammation, by activating important pathways like JAK/Stat, Ras, ERK and Akt pathways[17] [18].

PI3K-Akt signaling pathway play important roles in apoptosis, autophagy, metabolism, cell growth and differentiation. The expression of FoxP3 by inhibiting the activation of transcription factor Forkhead-O3a (Foxo1-3a) is negatively regulated by this pathway[19]. The FoxP3+Treg cells activation which will assist to set up a new target for the involvement of TB immunotherapy molecules as part of the immune-escape mechanism to provide a theoretical basis is inhibited by M.tuberculosis[20]. In the case of latent infection to TB disease few additional pathways are involved like Regulation of Actin cytoskeleton during MTB infection, to maintain the stability of the cytoskeleton, macrophages cells themselves are also trying to regulate cytoskeletal associated proteins[21]. Thyroid hormones (hormones, T4 and T3) are made by thyroid gland which is essential for the regulation of metabolic processes throughout the body.

The regulation of autophagy is important for host in response to invading mycobacteria, the naïve host defence system recognizes pathogen motifs through innate receptors but also produces appropriate effector proteins, including cytokines. These innate signals regulate autophagic pathways by up regulation of genes of this pathway during infection[22]. Recently, a study reports that the enzyme Msm_ACTase (from bacteria) aids in scavenging increased amount of H_2_O_2_ due to over expression of genes involved in peroxisome pathway which gives insight to a new mechanism how the pathogen surpass the host defense in Mtb infection[23]. Moreover, Macrophages are the main effector cells responsible for killing pathogen (MTB) via different mechanisms, including apoptosis but important genes in this pathway were down regulated by MTB that slow down or stop the apoptotic process[24]. Most of the MicroRNAs that were found deregulated in cancer are effected by Mtb in a similar way while its infestation like deletion, mutation, and epigenetic silencing[25].

In the field of pharmacogenomics, to understand the regulation of disease-network is the great application in the drug discovery. We have tried to build the networks emphasis on genes that are regulated by network and this constructed network of classified genes from various stages of TB shows hierarchical nature, that indicate the networks have system level organization including modules/sub-modules which are interconnected. Since the network’s nature is hierarchical, its synchronisation confirms several important functional regulations of the network, but individual gene-activities are not so important. In our six networks (including UP & Down regulated genes), a total of 29 key regulators were identified by affecting motifs and module regulation, showing their biological important and serve as the foundation of network activities and their regulations and could be a most probable target gene for disease control. We have noticed that in normal to latent infection two genes TNF and IL1B are present (down regulated) in most of the important pathways, from literature we found that TNF and IL1 (A & B) are key mediators present in severe inflammatory diseases. However, both TNF and IL-1 receptor pathways are essential for the control of Mycobacterium tuberculosis infection, and it is critical to assess the respective role of IL1A, IL1B, and TNF[26] [27]. Beside of CCL4 expression is high in late phase of the active disease but lower levels during early infection[28]. It has been currently reported that AREG play a central role in orchestrating both host resistance and tolerance mechanisms. Although AREG is known as epithelial cell-derived factor and recent studies show that AREG can be expressed by multiple populations of activated immune cells in inflammatory conditions[29]. In Host immune system, NFKB1 plays a major role in the activation of immune cells by upregulating the expression of many cytokines essential to the immune responses[30] and gene CD44 play important role in innate and adaptive immune responses, in the acute inflammatory response to both infectious and sterile stimuli and during infection CD44 may influence host defence by affecting phagocytosis[31] but in our study, we found that NFKB1 and CD44 are down regulated that means, MTB is burglarize and seizing the host immune system by slow down the expression of NFKB1. It has been reported that MET (Hepatocyte growth factor receptor) regulate various functions of immune cells, including differentiation and maturation, cytokine production, cellular migration and adhesion, and T cell effector function and HGF exerts anti-inflammatory activities through MET signaling[32]. A group of researchers has elucidated that PLAUR domain containing 8 (Lypd8) inhibits bacterial invasion of colonic epithelia[33]). The gene TRAF1 has diverse biological functions, acting through direct or indirect interactions with multiple tumor necrosis factor receptor (TNFR) and intracellular proteins. Several studies have shown that TRAF1 might exert an antiapoptotic role in lymphoma cells via regulation of the activation of NF-?B[34]. It has been shown that the gene TNFAIP3 (A20) is a cytoplasmic zinc-finger protein that is induced under inflammatory conditions and acts as a negative-feedback regulator of NF-κB activation in response to multiple stimuli[35]. It has been shown that IRAK2 and its genetic variants as critical factors and potentially novel biomarkers for human antiviral innate immunity[36] and MAP3K8 is a serine-threonine kinase has a critical function in integrating host immune responses to complex pathogens[37]. The down regulation of these genes has allowed the process of host defense to cease and give time to bacteria for progression. Besides of these down regulated genes few genes are up regulated as Host responses against pathogen e.g., NLRC4 gene is associated with inflammasome signaling, Its up regulation means Inflammasome activation (play an important role in host defence against Mtb) not only leads to cytokine secretion but may also cause pyroptosis[38] [39] and NDUFS8 genes also play important role in host immunity by increase expression level[40]. It has been reported that LRRK2 is involved in the ifn-γ response and host response to pathogens[41] and the genes DDX58 which interact with IRGM and promote its K63-linked polyubiquitination, indicating that IRGM is positioned at a nexus of various innate immunity[42] while SAMD9L and EIF2AK2 are play key roles in the innate immune responses to multiple stimuli, IF2AK2 is also involved in the regulation of signal transduction, apoptosis, cell proliferation and differentiation[43] [44] [45].

## Material and Method

The complete work flow of study is illustrated in Figure 7.

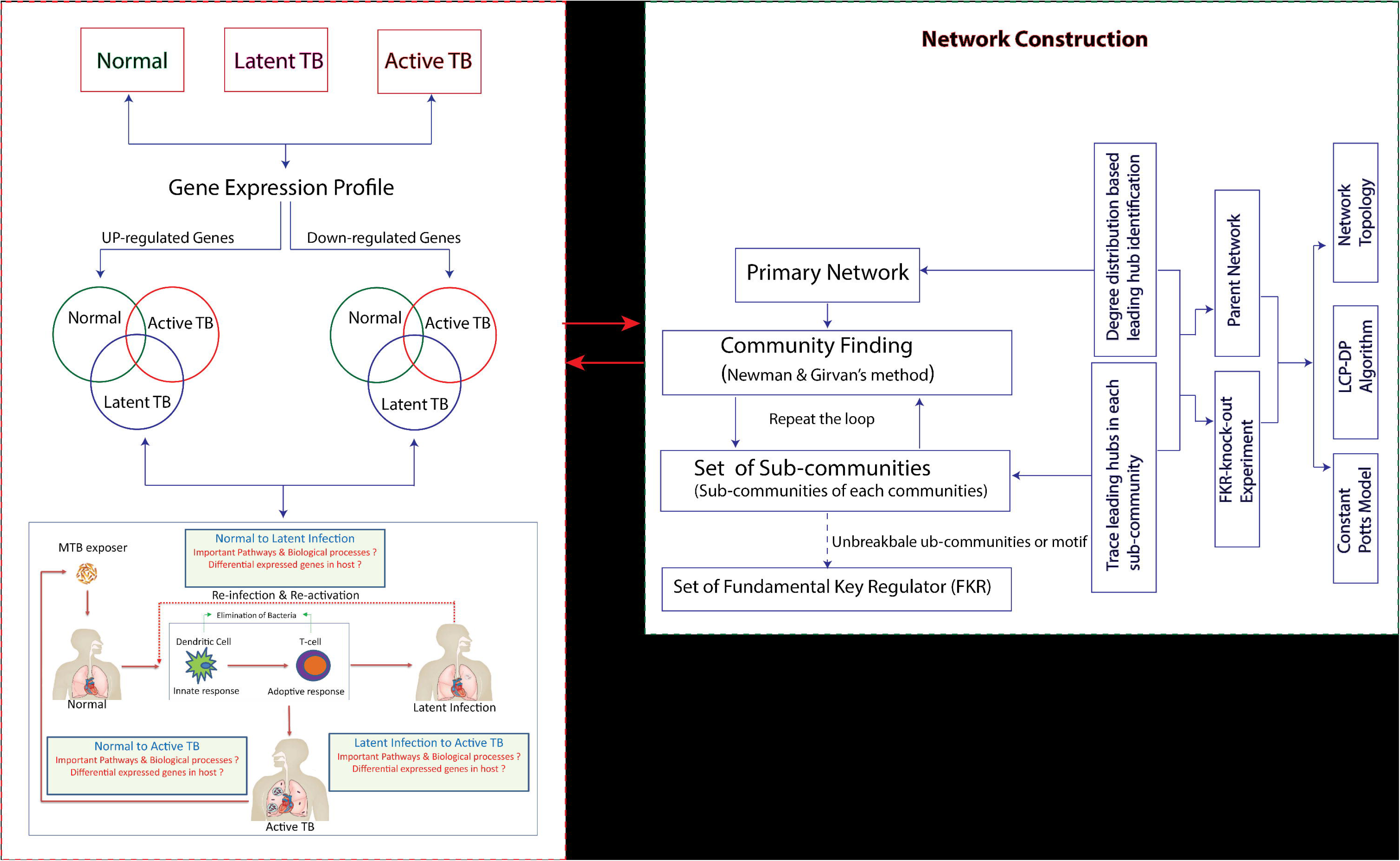

### 1. Inclusion Criteria for Differentially Expressed Genes

A set of 11 microarray data sets GSE54992[46], GSE52819[47], GSE64335[48], GSE11199[49], GSE98750[50], GSE78233[51], GSE57736[52], GSE16250[53], GSE27882[54] GSE57028[55], GSE83456[56] were selected from the NCBI GEO repository database (http://www.ncbi.nlm.nih.gov/geo/). It is currently largest and comprehensive public gene expression database. A total of 248 samples taken from the above GSE series, were divided into 3 clusters:(a) Healthy control (n=97), (b) Latent TB infection (n=12) and (c) active TB (n=139), shown in **Figure 8**. According to the corresponding correlation between the probe and gene from the data, the probe numbers of the expression profile were converted into the corresponding gene symbols using the Synergizer or DAVID [57] [58]. One gene corresponds with multiple probes, thus each gene has more than one expression value. Therefore, the average value was calculated and selected as the only representative value. Background correction and quartile data normalization were performed by the robust multiarray average in R affy package (Affymetrix). The limma package (Affymetrix)[59] was used to identify the DEGs among the Healthy control, Latent Tb and Active TB. The adjust P value < 0.05 and |logFC| ≥1.5 were set as the cut off criterion. We have used the BRCW (Bioinformatics & Research Computing website (http://jura.wi.mit.edu/bioc/tools/compare.php)) to select the DEGS which is same in at least two datasets of gene expression profile.

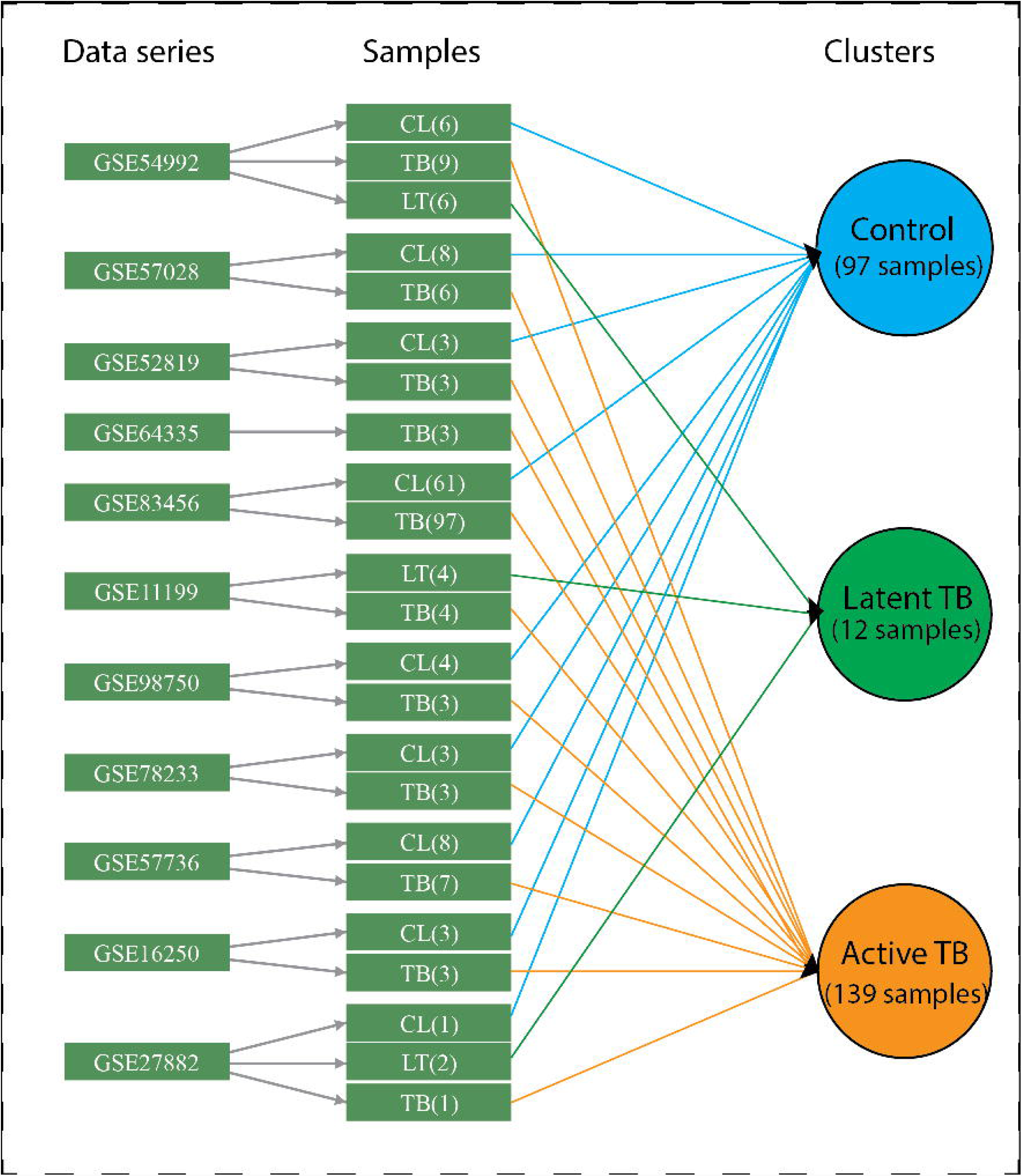

### 2. Gene Classification, Ontology and Pathway Analysis of DEGs

To know the significance of the identified DEGs, we have categorized them by GO-molecular function, GO-biological process and protein class using the Protein ANalysis THrough Evolutionary Relationships (PANTHER v.13.0) Classification System and analysis tools and DAVID (Database for Annotation Visualization and Integrated Discovery) an online software (https://david.ncifcrf.gov/) to enrich the given set of DEGs to possible GO terms [60] [61]. The PANTHER Overrepresentation study (Fisher’s Exact with FDR multiple test correction) was used to search the data against the PANTHER and GO databases and P-values were set according to Bonferroni correction.

The GO-analysis (Gene ontology) is the useful method For annotation of genes & its products and characterization of biological attributes for high-throughput genome or transcriptomes data[62]. The differentially expressed genes among ‘Active-TB’, ‘Latent TB’ and ‘Normal condition’ were over-represented in various GO classes. The gene ontology provides and visualize us the basic terms subdivided into three important categories namely BP (Biological process), MF (Molecular functions) and Biological pathways among those DEGs.

### 3. Gene-Transition among different stages of TB

In order to gets the behaviour of normal gene expression perturbation we tried a normal way for finding genes associated while moving from one stage to another. In all the transition we took into consideration has been provided by a list of UP and Down regulated genes, these gene(s) in both the cases (i.e. stage from which is transferred *to* the targeted stage) has its own meaning. This meaning to a gene(s) regulation gives us an opportunity to say something about the ongoing mechanobiology inside the cell. So, to observe these transitions we framed our study in such a way (based on the data we’ve got) that is to take every possible transition in view. These transitions are discussed in brief as follows. In this study, we made comparison of the gene-expression profiles among individuals with normal conditions, latent infection, active TB. Thus, we observed the expression of genes from normal to different stages of the TB and try to arrest those genes which play key role in progression of TB disease.

#### • Normal to Latent infection

In this section, we have taken those DEGs which are involved in between normal to latent TB condition. In LTBI, the bacteria remain inactive form for many years (years-decades) before transforming into TB disease. In this study, we have studied several biological processes, important pathways that lead to the latent infection for identification of latently infected individuals

#### • Normal to Active TB

In this section, we have taken those genes which are differentially expressed in TB disease condition as compared to normal and identified those immune process and pathways which are prominent in TB disease.

#### • Latent infection to Active

TB: In this section, we have taken those DEGs which is involved in between latent TB to active TB disease. The individual with Latent Tb infection eventually reactivates and becomes infectious, seriously influencing epidemiological situation. Mechanisms of MTB transition to dormancy and TB reactivation are inadequately understood, and biomarkers of latency remain largely mysterious [63].

### 4. Characterization of Topological Properties of Networks

The structural properties of complex networks are characterized through the behaviours of the topological parameters. The following topological properties of the networks were studied to learn the important behaviours of the network: *Degree distribution, Neighborhood connectivity, clustering co-efficient, Betweenness centrality, Closeness centrality* and *Eigenvector centrality.*

#### • Degree distribution

In a network, the degree **k** is a centrality measure that represents the number of links the node connects with other nodes. For a network defined by a graph ***G*= (*N,E*),** where **N** and **E** are number of nodes and edges respectively, the probability of degree distribution **(P(k))** of the network is the ratio of the number of nodes having degree to the network size;

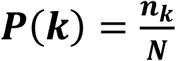

Where, **n_k_** is the number of nodes having degree **k** and **N** is the total number of nodes in the network. *P*(***k***) indicates the importance of hubs or modules in the network. It obeys power law **P(k) ~ k^-γ^** in scale-free and hierarchical networks depending on the value of **γ** which specifies the importance of hubs or modules in the network[64].

#### • Neighborhood connectivity

The number of neighbours of a node is considered as its connectivity. The neighborhood connectivity of a node n is defined as the average connectivity of all neighbors of n[65]. In the network **(CN(k))** Neighborhood connectivity is given by,

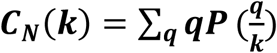

where, 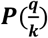 is the conditional probability that a link belonging to a node with connectivity **‘k’** points to a node with connectivity **‘q’.** The positive power dependence of **C_N_(k)** could be an indicator of assortativity in the network topology.

#### • Clustering co-efficient

This property of a network represents the measure of the interconnection of a node with its neighborhood node and strength of its interconnection. It is the ratio of the number of its nearest neighborhood edges e**_i_** to the total possible number of edges of degree **k_i_**. For an undirected network, clustering co-efficient **(C(k_i_))** of i^th^ node can be calculated by,

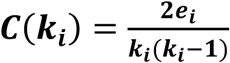

#### • Betweenness centrality

Betweenness centrality CB of a node represents the prominence of information flow in the network, and the extent to which the node has control over the other nodes in the network through communication[66], [67]. If **d_ij_ (v)** indicates the number of geodesic paths from node i to node j passing through node v, and dij indicates number of geodesic paths from node **i** to **j**, then betweenness centrality **(C_B_ (v))** of a node v can be calculated by,

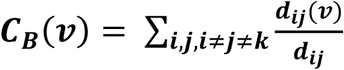

#### • Closeness centrality

Closeness centrality (C**_c_**) measures how fast information is spread from a node to other nodes accessible from it in the network[68]. The C**_c_** of a node i is the reciprocal of the mean geodesic distance between the node and all other nodes connected to it in the network, and is given by,

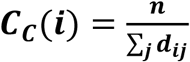

where **d_ij_** represents the geodesic path length from nodes **i** to **j**, and **n** is the total number of vertices in the graph reachable from node **i**.

#### • Eigenvector centrality

Eigenvector centrality of a node **i (C_E_ (i))** in a network is proportional to the sum of i’s neighbour centralities[69], and it is given by,

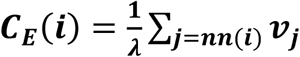

where**, nn(i)** indicates nearest neighbors of nodes i in the network. **λ** is eigenvalue of the eigenvector **v_i_** is given by, **Av_i_ = λv_i_** where, A is the adjacency matrix of the network (graph). The principal eigenvector of **A**, which corresponds to maximum eigenvalue **λ_max_**, is taken to have positive eigenvector centrality score. Eigenvector centrality can be used as an indicator of node’s spreading power in the network.

### 5. Community Analysis: Leading Eigen-vector method

In hierarchical network, to distinguish the nature of modular and its properties is important to explaining the predicting about the activities of network at various levels of hierarchy and access the organizing principle of the network. In our study, the Leading Eigen Vector method (LEV)[70], [71] was used to detect the communities in R from package ‘**igraph**’[72]. The LEV method is the most promising one for community detection as it calculates the Eigen value and exemplifying the significance for each link. To grub only motif, we detected modules from complete network and then sub-modules from the modules at each level of organization.

#### Modularity

Modularity determine how a network is divided in communities[73]. Modularity (Q) is expressed as follows,

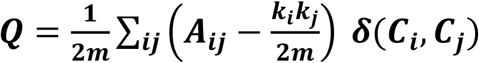

where m is the total number of edges in the community, A_ij_ is the adjacency matrix of size i × j, k represents degrees, and the δ function yields 1 if nodes i and j are in the same community.

### 6. Genes Tracing

To access the regulation of network, we first tried to find out the most influential nodes within the network. The gene tracing (up to motif level) was done purely on the appearance of the respective genes in various sub modules obtained from the clustering. Then these genes were used to get the picture of changes in the network-organization in their absence.

### 7. Hub Knock out Experiment

To know the changes of organization within the complex network in the absent of most influencing nodes, we must do removal of constructed rich-clubs or leading hubs (breaking monopoly) in the networks. We consecutively eliminate all the important hubs (one by one) from each network and measured the network properties of the reorganized network to characterize the regulating abilities of the hubs by calculating the degree of structural change due to their absent. The topological properties of the network were estimated using Network Analyzer, and for eigen value calculation, we used CytoNCA[74], plug-ins in Cytoscape for topological properties.

### 8. LCP-DP approach to estimate the network compactness

The representation of topological properties of a network in 2D parameter space of common neighbors **(CN)** index of interacting nodes and local community links (LCL) of each pair of interacting nodes in the network, and it provides information on number, size, and compactness of communities in a network, which can further be used as a measure of self-organization in the network[75]. The **CN** index between two nodes x and y can be calculated from the measure of overlapping between their sets of first-node-neighbors **S(x)** and **S(y)** given by, ***CN***= ***S(x***) ∩ ***S***(***y***). If there is significant amount of overlapping between the sets **S(x)** and **S(y)** (large value of **CN)**, the possible likelihood of interaction of these two nodes could happen and so an increase in CN represents the rise in compactness of the network, showing its faster information processing abilities. Further, the LCLs between the two nodes **x** and **y**, whose upper bound is defined by, 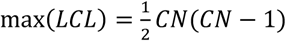 is the number of internal links in local-community (LC), which is strongly inter-linked group of nodes. Most probably these two nodes link together if **CN** of these two nodes are members of LC[76]. LCP-DP has been found to have a linear dependence between **CN** and 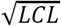. The LCP correlation (LCP-corr) is the Pearson correlation co-efficient between the variables CN and LCL defined by;

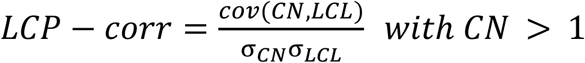

where *cov*(*CN*, ***LCL***) is the covariance among ***LCL*** & *CN* and σLCL & σCN are SD (standard deviations) of CN and LCL respectively.

### 9. Energy Distribution in the network: Hamiltonian energy calculation

The Hamiltonian energy (HE) is used to organize a network at a certain level by following the formalism of *Constant Potts Model*[77], [78]. The energy distribution at the global and modular level of a network is given by Hamiltonian energy (HE), which is in the self-1.organization of the system. Hamiltonian energy of a network or module or sub-module can be measured by,

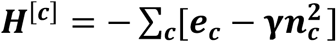

Where **e_c_** and **n_c_** are the number of edges and nodes in a community ‘**c**’ and ‘ **Y**’ is the resolution parameter acting as an edge density threshold which is set to be 0.5.

## Supporting information

## References

[1] S. Ahmad, “Pathogenesis, Immunology, and Diagnosis of Latent Mycobacterium tuberculosis Infection,” Clin. Dev. Immunol., vol. 2011, pp. 1–17, 2011.

[2] D. Schnappinger et al., “Transcriptional Adaptation of Mycobacterium tuberculosis within Macrophages: Insights into the Phagosomal Environment,” J. Exp. Med., vol. 198, no. 5, pp. 693–704, Sep. 2003.

[3] I. Lee, U. M. Blom, P. I. Wang, J. E. Shim, and E. M. Marcotte, “Prioritizing candidate disease genes by network-based boosting of genome-wide association data,” Genome Res., vol. 21, no. 7, pp. 1109–1121, Jul. 2011.

[4] A.-L. Barabási and Z. N. Oltvai, “Network biology: understanding the cell’s functional organization,” Nat. Rev. Genet., vol. 5, no. 2, pp. 101–113, Feb. 2004.

[5] R. Pastor-Satorras, A. Vázquez, and A. Vespignani, “Dynamical and Correlation Properties of the Internet,” Phys. Rev. Lett., vol. 87, no. 25, Nov. 2001.

[6] E. Ravasz and A.-L. Barabási, “Hierarchical organization in complex networks,” Phys. Rev. E, vol. 67, no. 2, Feb. 2003.

[7] A. Clauset, C. R. Shalizi, and M. E. J. Newman, “Power-Law Distributions in Empirical Data,” SIAM Rev., vol. 51, no. 4, pp. 661–703, Nov. 2009.

[8] Z.-X. Huang, H.-Y. Tian, Z.-F. Hu, Y.-B. Zhou, J. Zhao, and K.-T. Yao, “GenCLiP: a software program for clustering gene lists by literature profiling and constructing gene co-occurrence networks related to custom keywords,” BMC Bioinformatics, vol. 9, no. 1, p. 308, 2008.

[9] J.-H. Wang et al., “GenCLiP 2.0: a web server for functional clustering of genes and construction of molecular networks based on free terms,” Bioinformatics, vol. 30, no. 17, pp. 2534–2536, Sep. 2014.

[10] Z. Yang et al., “How dormant is Mycobacterium tuberculosis during latency? A study integrating genomics and molecular epidemiology,” Infect. Genet. Evol. J. Mol. Epidemiol. Evol. Genet. Infect. Dis., vol. 11, no. 5, pp. 1164–1167, Jul. 2011.

[11] S. D. Lawn and A. I. Zumla, “Tuberculosis,” Lancet Lond. Engl., vol. 378, no. 9785, pp. 57–72, Jul. 2011.

[12] M. Faridgohar and H. Nikoueinejad, “New findings of Toll-like receptors involved in Mycobacterium tuberculosis infection,” Pathog. Glob. Health, vol. 111, no. 5, pp. 256–264, Jul. 2017.

[13] M. Fallahi-Sichani, D. E. Kirschner, and J. J. Linderman, “NF-?B Signaling Dynamics Play a Key Role in Infection Control in Tuberculosis,” Front. Physiol., vol. 3, p. 170, 2012.

[14] K. Ozaki and W. J. Leonard, “Cytokine and cytokine receptor pleiotropy and redundancy,” J. Biol. Chem., vol. 277, no. 33, pp. 29355–29358, Aug. 2002.

[15] C.-H. Song et al., “Role of mitogen-activated protein kinase pathways in the production of tumor necrosis factor-alpha, interleukin-10, and monocyte chemotactic protein-1 by Mycobacterium tuberculosis H37Rv-infected human monocytes,” J. Clin. Immunol., vol. 23, no. 3, pp. 194–201, May 2003.

[16] G. Sabio and R. J. Davis, “TNF and MAP kinase signalling pathways,” Semin. Immunol., vol. 26, no. 3, pp. 237–245, Jun. 2014.

[17] A. Zlotnik and O. Yoshie, “Chemokines: a new classification system and their role in immunity,” Immunity, vol. 12, no. 2, pp. 121–127, Feb. 2000.

[18] A. Bajetto, R. Bonavia, S. Barbero, T. Florio, and G. Schettini, “Chemokines and Their Receptors in the Central Nervous System,” Front. Neuroendocrinol., vol. 22, no. 3, pp. 147–184, Jul. 2001.

[19] X. Zhang et al., “Inhibition of the PI3K-Akt-mTOR signaling pathway in T lymphocytes in patients with active tuberculosis,” Int. J. Infect. Dis., vol. 59, pp. 110–117, Jun. 2017.

[20] J. P. Scott-Browne et al., “Expansion and function of Foxp3-expressing T regulatory cells during tuberculosis,” J. Exp. Med., vol. 204, no. 9, pp. 2159–2169, Sep. 2007.

[21] J. Wang et al., “The fibroblast growth factor-2 arrests Mycobacterium avium sp. paratuberculosis growth and immunomodulates host response in macrophages,” Tuberc. Edinb. Scotl., vol. 95, no. 4, pp. 505–514, Jul. 2015.

[22] E.-K. Jo, “Autophagy as an innate defense against mycobacteria,” Pathog. Dis., vol. 67, no. 2, pp. 108–118, Mar. 2013.

[23] G. Ganguli, “Mycobacterium tuberculosis Acetyltransferase Reduces the Oxidative Stress Response Through Expression of Peroxisomal Membrane Transporter Protein,” Open Forum Infect. Dis., vol. 2, no. suppl_1, 2015.

[24] F. Behler et al., “Macrophage-Inducible C-Type Lectin Mincle-Expressing Dendritic Cells Contribute to Control of Splenic Mycobacterium bovis BCG Infection in Mice,” Infect. Immun., vol. 83, no. 1, pp. 184–196, Jan. 2015.

[25] R. Garzon, G. A. Calin, and C. M. Croce, “MicroRNAs in Cancer,” Annu. Rev. Med., vol. 60, pp. 167–179, 2009.

[26] M.-L. Bourigault et al., “Relative contribution of IL-1α, IL-1β and TNF to the host response to Mycobacterium tuberculosis and attenuated M. bovis BCG: IL-1α, IL-1β versus TNF in Host Response to M. tuberculosis,” Immun. Inflamm. Dis., vol. 1, no. 1, pp. 47–62, Oct. 2013.

[27] Y. V. N. Cavalcanti, M. C. A. Brelaz, J. K. de A. Lemoine Neves, J. C. Ferraz, and V. R. A. Pereira, “Role of TNF-Alpha, IFN-Gamma, and IL-10 in the Development of Pulmonary Tuberculosis,” Pulm. Med., vol. 2012, pp. 1–10, 2012.

[28] J. F. Rangel-Santiago et al., “A novel role of Yin-Yang-1 in pulmonary tuberculosis through the regulation of the chemokine CCL4,” Tuberculosis, vol. 96, pp. 87–95, Jan. 2016.

[29] D. M. W. Zaiss, W. C. Gause, L. C. Osborne, and D. Artis, “Emerging Functions of Amphiregulin in Orchestrating Immunity, Inflammation, and Tissue Repair,” Immunity, vol. 42, no. 2, pp. 216–226, Feb. 2015.

[30] S. Ghosh, M. J. May, and E. B. Kopp, “NF-?B AND REL PROTEINS: Evolutionarily Conserved Mediators of Immune Responses,” Annu. Rev. Immunol., vol. 16, no. 1, pp. 225–260, Apr. 1998.

[31] G. J. W. van der Windt et al., “CD44 Deficiency Is Associated with Increased Bacterial Clearance but Enhanced Lung Inflammation During Gram-Negative Pneumonia,” Am. J. Pathol., vol. 177, no. 5, pp. 2483–2494, Nov. 2010.

[32] Z. Sagi and T. Hieronymus, “The Impact of the Epithelial–Mesenchymal Transition Regulator Hepatocyte Growth Factor Receptor/Met on Skin Immunity by Modulating Langerhans Cell Migration,” Front. Immunol., vol. 9, Mar. 2018.

[33] R. Okumura et al., “Lypd8 promotes the segregation of flagellated microbiota and colonic epithelia,” Nature, vol. 532, no. 7597, pp. 117–121, Mar. 2016.

[34] X.-K. Wan et al., “Helicobacter pylori inhibits the cleavage of TRAF1 via a CagA-dependent mechanism,” World J. Gastroenterol., vol. 22, no. 48, p. 10566, 2016.

[35] L. Vereecke, R. Beyaert, and G. van Loo, “Genetic relationships between A20/TNFAIP3, chronic inflammation and autoimmune disease,” Biochem. Soc. Trans., vol. 39, no. 4, pp. 1086–1091, Aug. 2011.

[36] H. Wang et al., “A frequent hypofunctional IRAK2 variant is associated with reduced spontaneous hepatitis C virus clearance: WANG, EL MAADIDI, ET AL.,” Hepatology, vol. 62, no. 5, pp. 1375–1387, Nov. 2015.

[37] L. A. Mielke et al., “Tumor Progression Locus 2 (Map3k8) Is Critical for Host Defense against Listeria monocytogenes and IL-1 Production,” J. Immunol., vol. 183, no. 12, pp. 7984–7993, Dec. 2009.

[38] J. P.-Y. Ting, S. B. Willingham, and D. T. Bergstralh, “NLRs at the intersection of cell death and immunity,” Nat. Rev. Immunol., vol. 8, no. 5, pp. 372–379, May 2008.

[39] T. Bergsbaken, S. L. Fink, and B. T. Cookson, “Pyroptosis: host cell death and inflammation,” Nat. Rev. Microbiol., vol. 7, no. 2, pp. 99–109, Feb. 2009.

[40] B. K. Lohman, N. C. Steinel, J. N. Weber, and D. I. Bolnick, “Gene Expression Contributes to the Recent Evolution of Host Resistance in a Model Host Parasite System,” Front. Immunol., vol. 8, Sep. 2017.

[41] A. Gardet et al., “LRRK2 Is Involved in the IFN-Response and Host Response to Pathogens,” J. Immunol., vol. 185, no. 9, pp. 5577–5585, Nov. 2010.

[42] S. Chauhan, M. A. Mandell, and V. Deretic, “Mechanism of action of the tuberculosis and Crohn disease risk factor IRGM in autophagy,” Autophagy, vol. 12, no. 2, pp. 429–431, Feb. 2016.

[43] A. Lemos de Matos, J. Liu, G. McFadden, and P. J. Esteves, “Evolution and divergence of the mammalian SAMD9/SAMD9L gene family,” BMC Evol. Biol., vol. 13, no. 1, p. 121, 2013.

[44] K. Onomoto et al., “Critical Role of an Antiviral Stress Granule Containing RIG-I and PKR in Viral Detection and Innate Immunity,” PLoS ONE, vol. 7, no. 8, p. e43031, Aug. 2012.

[45] L. C. Reineke and R. E. Lloyd, “The Stress Granule Protein G3BP1 Recruits Protein Kinase R To Promote Multiple Innate Immune Antiviral Responses,” J. Virol., vol. 89, no. 5, pp. 2575–2589, Mar. 2015.

[46] Y. Cai et al., “Increased complement C1q level marks active disease in human tuberculosis,” PloS One, vol. 9, no. 3, p. e92340, 2014.

[47] M. Verway et al., “Vitamin D induces interleukin-1β expression: paracrine macrophage epithelial signaling controls M. tuberculosis infection,” PLoS Pathog., vol. 9, no. 6, p. e1003407, 2013.

[48] M. Bouttier et al., “Alu repeats as transcriptional regulatory platforms in macrophage responses to M. tuberculosis infection,” Nucleic Acids Res., vol. 44, no. 22, pp. 10571–10587, Dec. 2016.

[49] N. T. T. Thuong et al., “Identification of tuberculosis susceptibility genes with human macrophage gene expression profiles,” PLoS Pathog., vol. 4, no. 12, p. e1000229, Dec. 2008.

[50] L. Lavalett, H. Rodriguez, H. Ortega, W. Sadee, L. S. Schlesinger, and L. F. Barrera, “Alveolar macrophages from tuberculosis patients display an altered inflammatory gene expression profile,” Tuberc. Edinb. Scotl., vol. 107, pp. 156–167, 2017.

[51] C. Y. Cheng et al., “Host sirtuin 1 regulates mycobacterial immunopathogenesis and represents a therapeutic target against tuberculosis,” Sci. Immunol., vol. 2, no. 9, Mar. 2017.

[52] J. M. Guerra-Laso, S. Raposo-García, S. García-García, C. Diez-Tascón, and O. M. Rivero-Lezcano, “Microarray analysis of Mycobacterium tuberculosis-infected monocytes reveals IL26 as a new candidate gene for tuberculosis susceptibility,” Immunology, vol. 144, no. 2, pp. 291–301, Feb. 2015.

[53] N. Reyes, A. Bettin, I. Reyes, and J. Geliebter, “Microarray analysis of the in vitro granulomatous response to Mycobacterium tuberculosis H37Ra,” Colomb. Medica Cali Colomb., vol. 46, no. 1, pp. 26–32, Mar. 2015.

[54] K. Liu et al., “Increased levels of BAFF and APRIL related to human active pulmonary tuberculosis,” PloS One, vol. 7, no. 6, p. e38429, 2012.

[55] H. Salamon et al., “Cutting edge: Vitamin D regulates lipid metabolism in Mycobacterium tuberculosis infection,” J. Immunol. Baltim. Md 1950, vol. 193, no. 1, pp. 30–34, Jul. 2014.

[56] S. Blankley et al., “The Transcriptional Signature of Active Tuberculosis Reflects Symptom Status in Extra-Pulmonary and Pulmonary Tuberculosis,” PloS One, vol. 11, no. 10, p. e0162220, 2016.

[57] G. F. Berriz and F. P. Roth, “The Synergizer service for translating gene, protein and other biological identifiers,” Bioinforma. Oxf. Engl., vol. 24, no. 19, pp. 2272–2273, Oct. 2008.

[58] D. W. Huang et al., “The DAVID Gene Functional Classification Tool: a novel biological module-centric algorithm to functionally analyze large gene lists,” Genome Biol., vol. 8, no. 9, p. R183, 2007.

[59] G. K. Smyth, “Linear models and empirical bayes methods for assessing differential expression in microarray experiments,” Stat. Appl. Genet. Mol. Biol., vol. 3, p. Article3, 2004.

[60] H. Mi, A. Muruganujan, and P. D. Thomas, “PANTHER in 2013: modeling the evolution of gene function, and other gene attributes, in the context of phylogenetic trees,” Nucleic Acids Res., vol. 41, no. Database issue, pp. D377-386, Jan. 2013.

[61] D. W. Huang, B. T. Sherman, and R. A. Lempicki, “Systematic and integrative analysis of large gene lists using DAVID bioinformatics resources,” Nat. Protoc., vol. 4, no. 1, pp. 44–57, 2009.

[62] M. Ashburner et al., “Gene ontology: tool for the unification of biology. The Gene Ontology Consortium,” Nat. Genet., vol. 25, no. 1, pp. 25–29, May 2000.

[63] T. Kondratieva, T. Azhikina, B. Nikonenko, A. Kaprelyants, and A. Apt, “Latent tuberculosis infection: what we know about its genetic control?,” Tuberc. Edinb. Scotl., vol. 94, no. 5, pp. 462–468, Sep. 2014.

[64] R. Albert and A.-L. Barabási, “Statistical mechanics of complex networks,” Rev. Mod. Phys., vol. 74, no. 1, pp. 47–97, Jan. 2002.

[65] S. Maslov, “Specificity and Stability in Topology of Protein Networks,” Science, vol. 296, no. 5569, pp. 910–913, May 2002.

[66] Ulrik Brandes, “A Faster Algorithm for Betweenness Centrality,” J. Math. Sociol., vol. 25, pp. 163--177, 2001.

[67] O. Mason and M. Verwoerd, “Graph theory and networks in Biology,” IET Syst. Biol., vol. 1, no. 2, pp. 89–119, Mar. 2007.

[68] H. M. Reza, A. Urano, N. Shimada, and K. Yasuda, “Sequential and combinatorial roles of maf family genes define proper lens development,” Mol Vis, vol. 13, pp. 18–30, 2007.

[69] P. Bonacich, “Power and Centrality: A Family of Measures,” Am. J. Sociol., vol. 92, no. 5, pp. 1170–1182, Mar. 1987.

[70] M. E. J. Newman, “Finding community structure in networks using the eigenvectors of matrices,” Phys. Rev. E, vol. 74, no. 3, Sep. 2006.

[71] M. E. J. Newman, “Modularity and community structure in networks,” Proc. Natl. Acad. Sci., vol. 103, no. 23, pp. 8577–8582, Jun. 2006.

[72] Gabor Csardi and T. Nepusz, “The igraph software package for complex network research,” Inter J Comp Syst, vol. 1695, pp. 1–9, 2006.

[73] M. E. J. Newman and M. Girvan, “Finding and evaluating community structure in networks,” Phys. Rev. E Stat. Nonlin. Soft Matter Phys., vol. 69, no. 2 Pt 2, p. 026113, Feb. 2004.

[74] Y. Tang, M. Li, J. Wang, Y. Pan, and F.-X. Wu, “CytoNCA: a cytoscape plugin for centrality analysis and evaluation of protein interaction networks,” Biosystems, vol. 127, pp. 67–72, Jan. 2015.

[75] C. V. Cannistraci, G. Alanis-Lobato, and T. Ravasi, “From link-prediction in brain connectomes and protein interactomes to the local-community-paradigm in complex networks,” Sci. Rep., vol. 3, p. 1613, 2013.

[76] C. V. Cannistraci, G. Alanis-Lobato, and T. Ravasi, “From link-prediction in brain connectomes and protein interactomes to the local-community-paradigm in complex networks,” Sci. Rep., vol. 3, no. 1, Dec. 2013.

[77] V. A. Traag, p. Van Dooren, and Y. Nesterov, “Narrow scope for resolution-limit-free community detection,” Phys. Rev. E, vol. 84, no. 1, Jul. 2011.

[78] V. A. Traag, G. Krings, and P. Van Dooren, “Significant Scales in Community Structure,” Sci. Rep., vol. 3, no. 1, Dec. 2013.

