## Supplementary figures and images for "Stage specific classification of DEGs via statistical profiling and network analysis reveals potential biomarker associated with various stages of TB"

### Supplementary file 1

UP-regulated genes

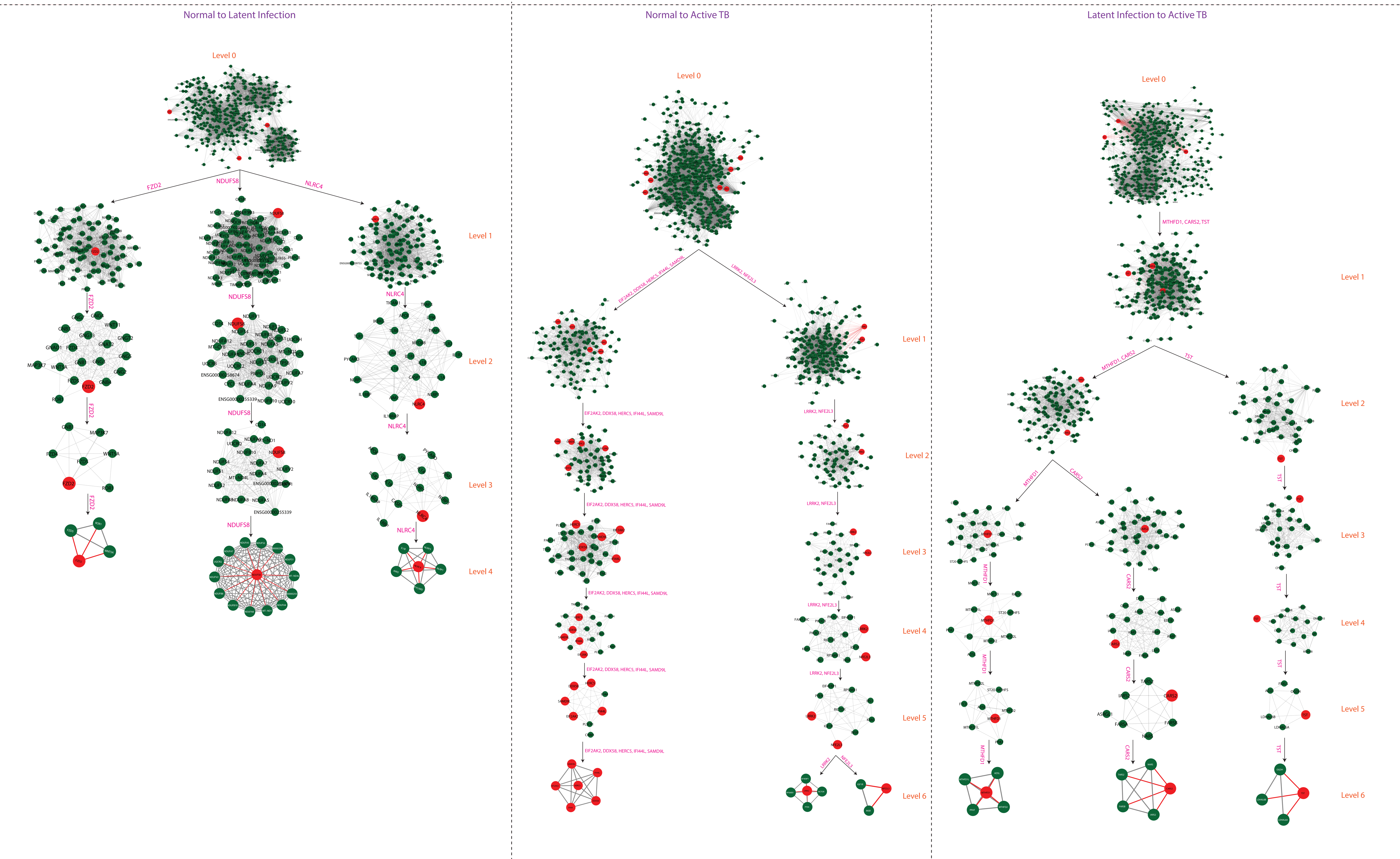

### Supplementary file 2

#### Down- regulated genes

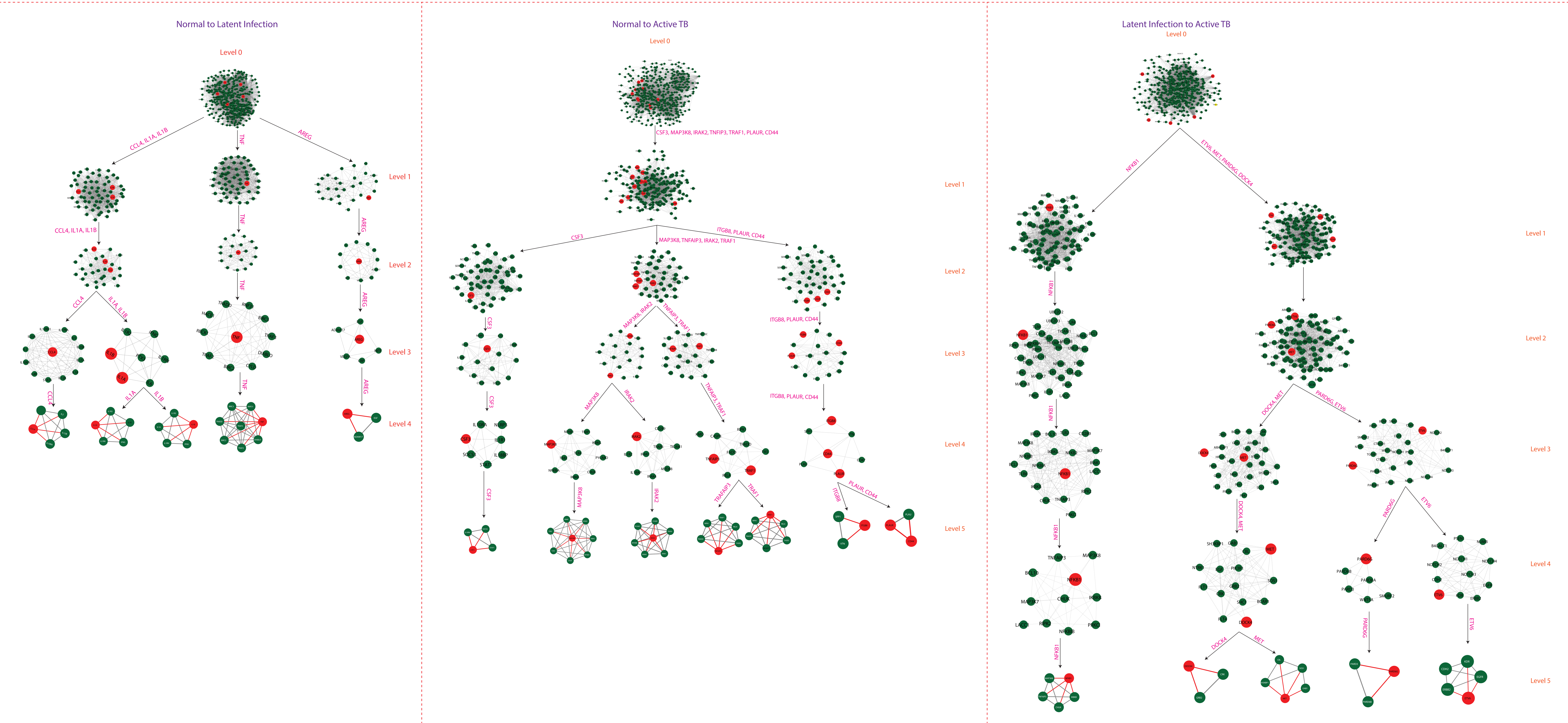
