## Supplementary material for "Stage specific classification of DEGs via statistical profiling and network analysis reveals potential biomarker associated with various stages of TB"

| Normal to Latent Infection | | |
|---|---|---|
| Gene ID | Uniprot ID | Protein name |
| FZD2 | Q14332 | Frizzled class receptor 2(FZD2) |
| NLRC4 | Q9NPP4 | NLR family CARD domain containing 4(NLRC4) |
| NDUFS8 | O00217 | NADH: ubiquinone oxidoreductase core subunit S8(NDUFS8) |
| *CCL4* | P13236 | C-C motif chemokine ligand 4 |
| *IL1B* | P01584 | Interleukin 1 beta(IL1B) |
| *IL1A* | P01583 | Interleukin 1 alpha(IL1A) |
| *TNF* | P01375 | Tumor necrosis factor(TNF) |
| *AREG* | P15514 | Amphiregulin(AREG) |
| **Normal to Active TB** | | |
| *EIF2AK2* | P19525 | Eukaryotic translation initiation factor 2 alpha kinase 2(EIF2AK2) |
| *SAMD9L* | Q8IVG5 | Sterile alpha motif domain containing 9 like(SAMD9L) |
| *IFI44L* | Q53G44 | Interferon induced protein 44 like(IFI44L) |
| *DDX58* | O95786 | DExD/H-box helicase 58 |
| *IFI44* | Q8TCB0 | Interferon induced protein 44(IFI44) |
| *HERC5* | Q9UII4 | HECT and RLD domain containing E3 ubiquitin protein ligase 5 |
| *NFE2L3* | Q9Y4A8 | Nuclear factor, erythroid 2 like 3(NFE2L3) |
| *LRRK2* | Q5S007 | Leucine rich repeat kinase 2(LRRK2) |
| *CSF3* | P09919 | Colony stimulating factor 3(CSF3) |
| *MAP3K8* | P41279 | Mitogen-activated protein kinase kinase kinase 8(MAP3K8) |
| *IRAK2* | O43187 | Interleukin 1 receptor associated kinase 2(IRAK2) |
| *TNFAIP3* | P21580 | TNF alpha induced protein 3(TNFAIP3) |
| *TRAF1* | Q13077 | TNF receptor associated factor 1(TRAF1) |
| *PLAUR* | Q03405 | Plasminogen activator, urokinase receptor(PLAUR) |
| *CD44* | P16070 | CD44 antigen |
| **Latent infection to Active TB** | | |
| *TST* | Q16762 | Thiosulfate sulfurtransferase(TST) |
| *MTHFD1* | P11586 | Methylenetetrahydrofolate dehydrogenase, cyclohydrolase and formyltetrahydrofolate synthetase 1(MTHFD1) |
| *CARS2* | Q9HA77 | Cysteinyl-tRNA synthetase 2, mitochondrial (putative)(CARS2) |
| *ETV6* | P41212 | ETS variant 6(ETV6) |
| *NFKB1* | P19838 | Nuclear factor kappa B subunit 1(NFKB1) |
| *MET* | P08581 | Hepatocyte growth factor receptor |

*UP-regulated

*Down-regulated

(A)

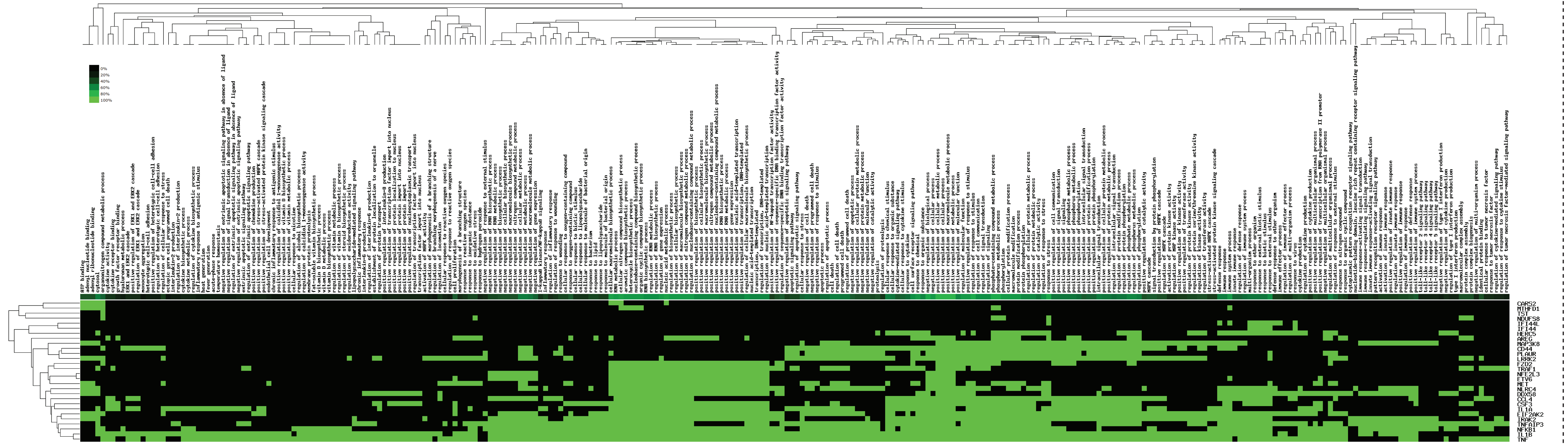

(B)
